# Semantic-Specific and Domain-General Mechanisms for Integration and Update of Contextual Information

**DOI:** 10.1101/2022.09.08.507135

**Authors:** Francesca M. Branzi, Matthew A. Lambon Ralph

## Abstract

Recent research has highlighted the importance of domain-general processes and brain regions for language and semantic cognition. Yet, this has been mainly observed in executively demanding tasks, leaving open the question of the contribution of domain-general processes to natural language and semantic cognition. Using fMRI, we investigated whether neural processes reflecting context integration and context update – two key aspects of naturalistic language and semantic processing – are domain-specific *versus* domain-general. Thus, we compared neural responses during integration of contextual information across semantic and non-semantic tasks. Whole-brain results revealed both shared (left posterior-dorsal inferior frontal gyrus, left posterior inferior temporal gyrus, and left dorsal angular gyrus/intraparietal sulcus) and distinct (left anterior-ventral inferior frontal gyrus, left anterior ventral angular gyrus, left posterior middle temporal gyrus for semantic control only) regions involved in context integration and update. Furthermore, data-driven functional connectivity analysis clustered domain-specific *versus* domain-general brain regions into distinct but interacting functional neural networks. These results provide a first characterization of the neural processes required for context-dependent integration during language processing along the domain-specificity dimension, and at the same time, they bring new insights on the role of left posterior lateral temporal cortex and left angular gyrus for semantic cognition.

## 1. Introduction

Recently, using combinations of functional magnetic resonance imaging (fMRI) and repetitive transcranial magnetic stimulation (rTMS) during naturalistic language processing, we established how the human brain supports buffering and integration of semantic information into a coherent contextual representation. That is, we revealed a distinction between the brain regions and neural networks that are crucial for the formation of a “semantic gestalt” (time-extended integration/formation of a semantic representation), *versus* those that are important for linking incoming cues about the current context (e.g., time and space cues) into a schema representation (Branzi, Humphreys, et al. 2020; Branzi, Pobric, et al. 2021).

Our previous research, however, left important issues unaddressed. For instance, it is unclear when and if the neural mechanisms that support buffering of information and its integration into a contextual representation reflect domain-specific or domain-general combinatorial operations. Furthermore, we still do not know very much about the semantic control processes required during formation of contextually integrated representations. For instance, are the semantic control demands required during integration of partially incoherent contexts (*hard* semantic integration) analogous to domain-general executive control processes? And, to what degree are the neural mechanisms, underpinning semantic control during context integration, specialized for this domain? The present fMRI study was designed to address these important questions.

Semantic cognition is supported by distributed neural network of fronto-parietal and temporal regions (Jefferies and Lambon Ralph 2006; Jefferies 2013; Canini et al. 2016; Lambon Ralph et al. 2017; Branzi, Martin, et al. 2021). Within this network, some brain regions have a key role in manipulating activation within the semantic representational system to generate semantic behaviours that are appropriate for the context in which they occur (Noonan, Jefferies, Visser and Lambon Ralph 2013; Lambon Ralph *et al*. 2017; Jackson 2020; Jackson et al. 2021). These brain regions include the inferior frontal gyrus (IFG) and the posterior lateral temporal cortex (pLTC) (particularly the posterior middle temporal gyrus – pMTG), which are strongly interlinked with functional and anatomical connections (Catani et al. 2005; Saur et al. 2008; Turken and Dronkers 2011) and are often co-activated for flexible and controlled retrieval of context or task-relevant semantic information (Jefferies and Lambon Ralph 2006; Whitney, Kirk, et al. 2011; Whitney et al. 2012; Noonan, Jefferies, Visser and Lambon Ralph 2013). Accordingly, we revealed that a discordant change in the semantic context or topic during naturalistic language processing induces strong activations in both IFG and middle pLTC/pMTG (Branzi, Humphreys, *et al*. 2020).

The left ventral angular gyrus (AG) is another key region for semantic cognition and control (Lambon Ralph *et al*. 2017; Jackson 2020). Differently from IFG and pLTC/pMTG, however, left ventral AG has been associated to a domain-general multimodal buffering function (Humphreys and Lambon Ralph 2015). That is, the ventral AG would operate as an online, dynamic buffer of any type of internal and external information that unfolds over time (Humphreys et al. 2019; Humphreys et al. 2021). Furthermore, the peak location for buffering information varies in a graded manner across the ventral AG, in line with the variations in long-range connections, such that anterior ventral AG is maximally recruited for buffering verbal/auditory information, core mid-AG for episodic information, and posterior ventral AG for visuo-spatial content. Consistent with this view, we recently demonstrated that ‘disrupting’ the left core-mid AG’s activity during buffering of contextual information impairs the encoding of contextually integrated representations (Branzi, Pobric, *et al*. 2021).

The left dorsal AG/intraparietal sulcus (dorsal AG/IPS) has been also associated to semantic cognition (Noonan, Jefferies, Visser and Lambon Ralph 2013; Lambon Ralph *et al*. 2017; Jackson 2020; Humphreys *et al*. 2021). However, differently from ventral AG, the dorsal AG/IPS receives connections from executively related brain regions, such as for instance the dorso-lateral prefrontal cortex, which are part of the ‘multi-demand-network’ (MDN), a set of brain regions activated across a broad range of executively demanding tasks (Duncan 2010; Fedorenko et al. 2013; Assem et al. 2020; Duncan et al. 2020). Accordingly, we have shown that dorsal AG/IPS is strongly recruited during integration of partially incoherent contexts (*hard* semantic integration) (Branzi, Humphreys, *et al*. 2020; Branzi, Pobric, *et al*. 2021).

There is a certain agreement that all the above-mentioned regions are important for semantic cognition. Yet, whether the regions implicated in integration of coherent and incoherent contexts, do so through domain-specific or domain-general processes remains unclear. Some have proposed that specific sub-regions of left IFG, left pLTC and left AG might support domain-general control processes (Hodgson et al. 2021). These claims are motivated by the observation of some overlap between the neural network for semantic control and the MDN (Geranmayeh et al. 2014; Humphreys and Lambon Ralph 2015; Hodgson *et al*. 2021). However, a minority of studies have directly compared semantic *versus* non-semantic tasks tapping similar cognitive operations (e.g., context integration), and those that have obtained mixed results (Humphreys et al. 2015; Humphreys and Lambon Ralph 2017; Humphreys *et al*. 2019; Quillen et al. 2021). Perhaps more problematic, the evidence of shared neural substrates between semantic and domain-general control processes is largely based on paradigms where typically little or no context information is provided during semantic processing, and/or where the main task is accompanied by a secondary task (Humphreys *et al*. 2015; Hodgson *et al*. 2021). As a result, such paradigms may indeed recruit domain-general control processes and the MDN, which is highly sensitive to task demands. However, this recruitment does not necessarily speak to the role of this network in core semantic operations, such as for instance formation and updating of a continuously evolving representation, i.e., the target of the present study.

A full characterization of the role of left IFG, pLTC and left AG for context integration and update requires considering their functional recruitment beyond the language/semantic domain. To this end, we conducted an fMRI study where we compared semantic *versus* non-semantic time-extended receptive tasks, made up of either two paragraphs of written text or number sequences, respectively (see **Figure 1** and Branzi *et al*., 2020 for a complete list of the verbal stimuli). In both tasks, there were three variations in respect to the relationship between the two paragraphs: (a) high congruency (HC condition) – the second paragraph continued with the same type of information, i.e., a coherent continuation of the same meaning/number sequence; (b) low congruency – where the second paragraph represented a change in the meaning or number sequence (LC condition); or (c) no context (NC condition) – the second paragraph was not preceded by any informative context (a different kind of material altogether). The comparison of HC and LC against NC condition allowed us to establish the brain regions important for integration of contextual information, i.e., how the brain builds an evolving representation. Instead, the comparison of HC against LC conditions allowed us to identify the brain regions involved in updating contextual information during integration. To address the key question of this study, for both contrasts, we examined the extent to which similar or dissimilar brain regions were engaged in semantic and non-semantic tasks.

**Figure 1.**
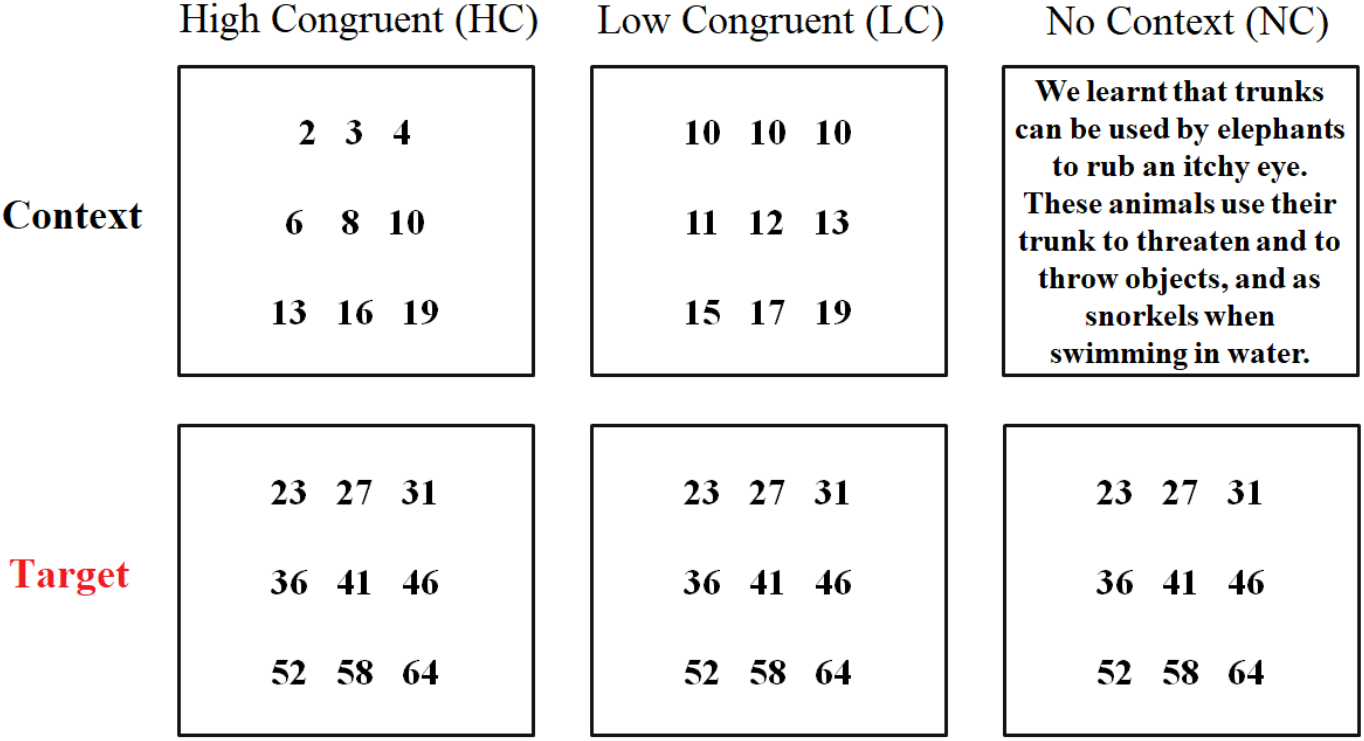
Example of trials in the non-semantic task, including HC, LC and NC conditions.

We expected both left IFG and pLTC to be implicated in the integration of contextual information (LC and HC > NC), especially when integration demands increase (LC > HC) (Noonan, Jefferies, Visser and Lambon Ralph 2013; Jackson 2020). However, we expected to observe a certain degree of functional heterogeneity within left IFG and pLTC. That is, the recruitment of different subregions depending on the type of stimuli/task (Hodgson *et al*. 2021). Specifically, for left IFG, we expected anterior-ventral portions (~Pars Orbitalis) to show domain-specific effects, i.e., to be positively engaged only in the semantic task (Hodgson *et al*. 2021). On the other hand, we expected more posterior-dorsal portions of left IFG (~Pars Triangularis and Pars Opercularis) to show domain-general effects (Badre and Wagner 2002; Dobbins and Wagner 2005; Branzi, Martin, et al. 2020).

For left pLTC, we expected left pMTG to be exclusively involved in semantic control (Hodgson *et al*. 2021), whilst the inferior portion (posterior inferior temporal gyrus - pITG) to show domain-general responses, in accord with evidence showing a strong link between neural activity in this region, the MDN and domain-general control processes (Assem *et al*. 2020; Hodgson *et al*. 2021).

We expected left AG to be involved in context integration (HC and LC > NC) (Branzi, Pobric, *et al*. 2021; Humphreys *et al*. 2021). However, recent explorations have shown that the AG is not a homogenous region, with some sub-regions that seems to be implicated in domain-specific and others in domain-general processes. Specifically, dorsal AG/IPS is active across a large variety of tasks and cognitive activities, especially when task or conditions demands increase (Noonan, Jefferies, Visser and Lambon Ralph 2013; Humphreys and Lambon Ralph 2017; Humphreys *et al*. 2019). Accordingly, we expected left dorsal AG/IPS to be engaged in both tasks during integration of contextual information, and more strongly for LC as compared to HC conditions (Noonan, Jefferies, Visser and Lambon Ralph 2013). In contrast, the left anterior ventral AG shows a preference for verbal stimuli, such as sentences or narratives, especially when task demands decrease (Humphreys and Lambon Ralph 2015; Branzi, Humphreys, *et al*. 2020). However, the profile of activation/deactivation of the ventral AG in this study is hard to predict from the current literature. In fact, on the one hand, some results have shown that the ventral AG is positively engaged for processing of verbal/semantic stimuli, but deactivated for numerical stimuli (Humphreys *et al*. 2019). On the other hand, other results suggest that this region might be positively engaged during numerical and verbal/semantic processing alike, especially when control demands decrease (Humphreys and Lambon Ralph 2015).

To fully characterize the profile of engagement of different IFG, pLTC and AG subregions during semantic and non-semantic tasks, we also employed multivariate functional connectivity analysis, and particularly independent component analysis (ICA). This data-driven approach allowed us to (1) measure if the above-mentioned brain regions exhibited a “yoked” activation, i.e., if they formed coherent functional networks; (2) establish if the networks that support contextual integration and update show domain-specific or domain-general patterns of engagement; and finally, (3) examine the extent to which interactions between domain-specific and domain-general networks are modulated by the task domain.

Based on previous evidence, we expected to observe both domain-general and domain-specific networks (Geranmayeh *et al*. 2014; Humphreys *et al*. 2019). In detail, we expected a fronto-parietal network including domain-general brain regions (i.e., posterior-dorsal portions of IFG, left dorsal AG/IPS and inferior pLTC/pITG) to be similarly involved in the two tasks, and to reflect context integration processes (HC & LC > NC). We expected domain-specific networks to include domain-specific subregions, such as for instance the ATL for the semantic task. Since the language/semantic network – but not the MDN – may track semantic content specifically (Diachek et al. 2020), we expected domain-specific networks specifically to support the update of the ongoing representation (LC > HC). Finally, we also expected some spatial overlap and interaction between the domain-general network for context integration and domain-specific networks for context update. Specifically, we expected spatial overlap in brain regions reflecting domain-general neural responses and inter-networks functional interactions to be modulated by the type of task or stimuli employed.

## 2. Materials and Methods

### 2.1. Participants

All participants were native English speakers with no history of neurological or psychiatric disorders, and normal or corrected-to-normal vision. Twenty-four volunteers took part in the semantic task (average age=22, SD=2; N female=18) (Branzi, Humphreys, *et al*. 2020). As a result of technical issues during the scanning session, only data from 22 participants study (average age=22, SD=2; N female=18) were usable for fMRI data analyses. Twenty-four volunteers took part in the non-semantic task (average age=22, SD=2; N female=19). Both experiments were approved by the local ethics committee. Written informed consent was obtained from all participants.

### 2.2. Stimuli

#### Semantic task

A total of 40 narrative pairs were created. For each narrative pair, the same second paragraph (target) was preceded by different first paragraphs (contexts) that could be either high-congruent (i.e., HC) or low-congruent (i.e., LC) with the target in terms of meaning. Both HC and LC context paragraphs could be integrated with the final target paragraphs, though a reworking of the evolving semantic context was required after LC contexts only, because of a shift in the semantic context (comprehensive list of the stimuli is reported in (Branzi, Humphreys, *et al*. 2020).

To confirm that HC and LC conditions differed in the semantic associative strength between contexts and targets, we quantified semantic relatedness between the contexts and targets, for both HC and LC conditions, in multiple ways. First, we employed Latent Semantic Analysis (LSA) to measure the semantic relationship between words based on the degree to which they are used in similar linguistic contexts. Thus, for each narrative pair, context and target paragraphs were converted in vectors of words that were successively compared, using a cosine similarity metric. From this comparison, an LSA value reflecting the associative strength between the context and target was obtained for both conditions. Results from LSA confirmed that semantic associative strength between the (same) target and the context was higher for HC (average score=0.4, SD=0.14) than LC conditions (average score=0.25, SD=0.09) [*t* (78) =-5.435,*p* < .001].

Second, as reported previously (Branzi, Humphreys, *et al*. 2020), we asked a group of independent participants to rate how semantically related contexts and targets were (0 to 5 scale). The results of this pre-experimental rating indicated that HC (average score=4.4, SD=0.4) and LC (average score=2.3, SD=0.4) conditions were different [*t* (9) =-10.626, *p* < .001]. Moreover, to ensure that participants could perceive the shift of semantic context during the study, at end of each narrative the question “Was there any change of semantic context between part 1 and part 2?” was posed. Only pairs of narratives on which at least the 90% of participants responded correctly to the questions were employed in the study.

Finally, another condition was included in the experimental design to measure the context integration processes. Specifically, in the no-context (NC) condition, the target (the same as in HC and LC conditions) was preceded by a string of numbers that could include from one to four-digit numbers.

#### Non-semantic Task

To compare the neural basis of context integration across semantic and non-semantic tasks, response times (RTs) were matched across the semantic and non-semantic domains. This was achieved by conducting a pilot study with 10 participants on preliminary versions of the non-semantic experiment outside the scanner. Based on these participants’ behavioural data, we iteratively adjusted several aspects of the stimuli of the non-semantic task until we obtained a final version in which conditions were matched to the semantic task in terms of RTs (Branzi, Humphreys, *et al*. 2020). Like the semantic task, a total of 40 stimuli pairs, each one composed by two “paragraphs”, were created. However, the content of each paragraph corresponded to numerical sequences. Thus, each paragraph contained a numerical sequence presented over three rows, with three numerical elements per row (see **Figure 1**). For each pair, the same second paragraph (target) was preceded by different first numerical paragraphs (contexts) that could be either high-congruent (i.e., HC) or low-congruent (i.e., LC) with the target paragraph’s numerical sequence. Both HC and LC context paragraphs could be associated with the target’s, though a change in the context was required after LC contexts only, because of a shift in the numerical sequence (**Figure 1**). Finally, as per the semantic task, another condition was included in the design to measure the context integration processes. Specifically, in the NC condition, the target paragraph (the same as that used in the HC and LC conditions) was preceded by a verbal paragraph taken from the semantic task (**Figure 1**).

### 4.3. Task procedures

#### Semantic task

There were 40 items per condition presented using an event-related design with the most efficient ordering of events determined using Optseq (http://www.freesurfer.net/optseq). Null time was intermixed between trials and varied between 2 and 12 seconds (s) (average=3.7, SD=2.8) during which a red fixation cross was presented. The red colour was used in order to mark the end of each trial (each trial composed by a context and a target). A black fixation cross was presented between contexts and targets and its duration varied between 0 and 6 sec (average=3, SD=1.6). Each context paragraph was presented for 9 sec followed by the target for 6 sec.

Participants were asked to read silently both contexts and targets that were displayed on the screen for 9 sec and 6 sec, respectively. Our volunteers were instructed to press a button when reaching the end of each paragraph (for both contexts and targets). The instruction emphasized speed, but also the need to understand the meaning of contexts and targets, since at the end of some of the trials, participants would be asked some questions about the content of the stimuli. We specified that, in order to perform this task, it would be necessary to process contexts and targets together. Hence, following 13% of the trials, a comprehension task was presented to ensure that participants were engaged in the task. When this happened, the target item was followed by a question displayed on the screen for 6 sec, and participants were asked to provide a response (true/false) via button press. A fixation cross, between the target and the question, was presented for a variable duration between 0 and 6 sec (average=3.5, SD=2.2). Before starting the experimental study, all participants signed an informed consent and were given written instructions. Then, they completed a practice session to familiarise with the task. The stimuli used in the practice session were different from those used in the experimental study.

#### Non-semantic Task

There were 40 items per condition that were presented with the same procedure as in the semantic task (see above). Thus, participants were asked to read silently both contexts and targets (displayed on the screen for 9 sec and 6 sec, respectively), process them as a unique numerical sequence, and to press a button when reaching the end of each paragraph (for both contexts and targets). However, to ensure that our volunteers were engaged in similar processing as in the semantic task (i.e., the same item-by-item and sentence-by-sentence integration processes), we also instructed participants to detect the combinatorial rule linking the first three numbers in the first row (e.g., incremental addition, see **Figure 1**) with the content of the other remaining rows.

Furthermore, we informed participants that sometimes the combinatorial rule would change, but that their task would nevertheless be the same, i.e., read the numbers presented in the context and target paragraphs silently, and detect the new combinatorial rule. The instruction emphasized speed, but also the need to process the information included in contexts and targets, since at the end of some trials participants would be asked questions on the content of the stimuli. As per the semantic task, following 13% of the trials, a comprehension task was presented to ensure that participants were engaged in the task. When this happened, the target item was followed by a question displayed on the screen for 6 sec, and participants were asked to provide a response (true/false) via button press. The questions pertained to the combinatorial rule(s) linking the number sequences, as well as the specific numbers presented in the context and target paragraphs. A fixation cross between the target and the question was presented for a variable duration between 0 and 6 sec (average=3.5, SD=2.2). Before starting the experimental study, all participants signed an informed consent and were given written instructions. Then, they completed a practice session to familiarize with the task. The stimuli used in the practice session were different from those used in the experimental study.

### 2.4. Task acquisition parameters

Images for both semantic and non-semantic tasks were acquired using a 3T Philips Achieva scanner using a dual gradient-echo sequence, which is known to have improved signal relative to conventional techniques, especially in areas associated with signal loss (Halai et al. 2014). Thus, 31 axial slices were collected using a TR=2 sec, TE=12 and 35 msec, flip angle = 95°, 80 × 79 matrix, with resolution 3 × 3 mm, slice thickness 4 mm. For each participant, 1492 volumes were acquired in total, collected in four runs of 746 sec each.

### 2.5. Data analysis

#### Behavioural data analyses

Since both tasks were reading tasks, accuracy measures could not be recorded. Instead, speed of reading (RTs) was analysed via a repeated-measures analysis of variance (ANOVA), with “Condition” as within-subjects factor with three levels (NC, LC and HC conditions) and “Task” (semantic task and non-semantic task) as a between-subject factor. To examine whether participants were similarly engaged in both tasks, we also computed an ANOVA including the percentage of given responses, with “Condition” as within-subjects factor with three levels (NC, LC and HC conditions) and “Task” (semantic task and non-semantic task) as a between-subject factor. Bonferroni correction for multiple comparisons was applied. Finally, correction for non-sphericity (Greenhouse–Geisser procedure) was applied to the degrees of freedom and *p* values associated with factors having more than two levels (i.e., Condition).

#### fMRI data analyses

##### Preprocessing

The dual-echo images were averaged. Data were analysed using SPM12. After motion-correction, images were co-registered to the participant’s T1 image. Spatial normalisation into MNI space was computed using DARTEL (Ashburner 2007), and the functional images were resampled to a 3 × 3 × 3mm voxel size and smoothed with an 8mm FWHM Gaussian kernel.

#### First level analysis

##### General Linear Modelling (GLM)

The data were filtered using a high-pass filter with a cut-off of 128s. For both datasets, we ran a GLM model in which, at the individual subject level, each condition of interest was modelled with a separate regressor (target NC, target LC and target HC) with time derivatives added, and events were convolved with the canonical hemodynamic response function, starting from the onset of the target paragraph. The context paragraphs (NC, LC and HC contexts) and the comprehension task were modelled as regressors. However, these data were not further analysed because they were not relevant for the scope of the present study. Each condition was modelled as a single event with a duration corresponding to 6 sec (target conditions and comprehension task trials) or 9 sec (context conditions). Motion parameters were entered into the model as covariates of no interest.

#### Second level analysis

##### Context integration and update in semantic and non-semantic tasks

To establish which brain regions were similarly engaged during contextual integration (HC & LC > NC) and update (LC > HC) of semantic *versus* non-semantic information, we conducted a flexible factorial 2 (semantic task, non-semantic task) × 3 (HC, LC, NC target condition) ANOVA, where subjects were treated as a random effect. The factor matrix included images derived from the first level analysis, relative to the NC, LC and HC target regressors from both semantic and non-semantic tasks. The main effects and interactions relative to the effects of interest were examined via whole-brain analyses (*F*-tests). These statistical maps were thresholded at *p* < .001 for voxel intensity, and *p* < .05 (family-wise error (FWE)- correction for multiple comparisons) for clusters. We also conducted a conjunction analysis to reveal which voxels showed, for each specified contrast of interest (i.e., HC & LC > NC and LC > HC), significant effects in both tasks. Finally, to investigate the direction of any potential difference between semantic and non-semantic tasks and conditions, as well as the engagement of these regions as compared to rest, the contrast estimates were also extracted and plotted via region of interest (ROI) analysis using spheres of 10mm radius. The data from ROI analysis were then analysed via ANOVAs, where “Task” was a between-subjects factor with two levels (semantic and non-semantic) and “Condition” a within-subjects factor with three levels (NC, LC and HC targets). Bonferroni correction for multiple comparisons was applied to assess statistically significant effects. Finally, correction for non-sphericity (Greenhouse–Geisser procedure) was applied to the degrees of freedom and *p* values associated with factors having more than two levels (i.e., Condition).

##### Task group spatial ICA in semantic and non-semantic tasks

Spatial ICA applied to fMRI data identifies temporally coherent networks by estimating maximally independent spatial sources, referred to as spatial maps, from their linearly mixed fMRI signals, referred to as time courses (Calhoun et al. 2001). We employed ICA to examine whether key brain areas revealed by univariate analyses were coupling with similar brain regions across semantic and non-semantic tasks to support context integration and update. Furthermore, with ICA we aimed to reveal whether functional interactions between domain-specific and domain-general networks were modulated by the type of stimuli (see Introduction).

The pre-processed fMRI data of both semantic and non-semantic tasks was analysed together in a group spatial ICA using the GIFT toolbox (http://mialab.mrn.org/software/gift) (Calhoun *et al*. 2001) to decompose the data into components. GIFT was used to concatenate the subjects’ data, and reduce the aggregated data set to the estimated number of dimensions using principal component analysis, followed by ICA using the Infomax algorithm (Bell and Sejnowski 1995). Subject-specific spatial maps and time courses were estimated using GICA back-reconstruction method based on principal component analysis compression and projection (Calhoun *et al*. 2001).

The number of independent components estimated within the data was 33. The estimation was achieved by using the Minimum Description Length criteria, first per each individual data-set and then computing the group mean. The obtained 33 independent components were inspected in order to exclude from the analysis artefactual and noise-related components. Similar to previous studies (Geranmayeh *et al*. 2014; Griffanti et al. 2017), the criterion for assigning components as artefact was based on the spatial maps attained as a result of the one sample *t*-tests (threshold for voxel-wise significance was set at *p* < .05, corrected for FWE). The spatial maps were visually compared with the SPM grey matter template. Only components that had the majority of activity within the grey matter were selected (N=26).

##### Establishing task-related functional networks in semantic and non-semantic task

The 26 independent components were labelled according to the resting state networks template provided in the GIFT toolbox. Then, a multiple regression analysis (implemented as “temporal sorting” function in GIFT) between independent component’s and task model’s time courses for each participant was conducted. This analysis allowed to identify which independent components reflected task-related functional networks in both tasks. In detail, for each participant the design matrix used for the GLM analysis, where rest periods were modelled implicitly as task baseline, was employed. For each independent component, the multiple regression analysis generated three beta weight values (one for each condition NC, LC, and HC) that were averaged across runs and participants. Beta weight values represent the correlations between time courses of the independent components and the canonical hemodynamic response model for each task condition. These values are thought to reflect engagement of the functional networks during specific task conditions (Xu et al. 2013).

After extracting the beta weight values for each independent component associated with each task-condition in each task, task-relatedness for each component was assessed. This was achieved by testing group means of averaged beta weight values for each task-condition against zero (one-sample *t*-tests, *p* < .05). Hence, a positive/negative beta weight, significantly different from zero, indicated an increase/decrease in activity of the independent component during a specific task condition relative to the baseline condition (i.e., rest). Having established the task-related functional networks, a repeated-measures ANOVA was used to assess the main differences between beta weight values across different task-conditions and tasks, for each network. Bonferroni correction for multiple comparisons was applied to assess statistically significant effects. Finally, correction for non-sphericity (Greenhouse–Geisser procedure) was applied to the degrees of freedom and *p* values associated with factors having more than two levels (i.e., Condition).

##### Functional network connectivity (FNC) analysis

To explore functional interactions between neural networks of interest (domain-specific and domain-general functional networks) we conducted a FNC analysis using the Mancovan toolbox in GIFT. Hence, FNC was estimated as the Pearson’s correlation coefficient between pairs of time courses (Jafri et al., 2008). Subject specific time courses were detrended and despiked based on the median absolute deviation as implemented in 3dDespike (http://afni.nimh.nih.gov/), then filtered using a fifth-order Butterworth low-pass filter with a high frequency cutoff of 0.15 Hz. Pairwise correlations were computed between time courses, resulting in a symmetric c1 × c1 correlation matrix for each subject. For all FNC analyses, correlations were transformed to z-scores using Fisher’s transformation, z = atanh(k), where k is the correlation between two component time courses. One sample t-tests (corrected for multiple comparisons at α = .01 significance level, using false discovery rate - FDR) were conducted on task-related functional networks to reveal the significance of pairwise correlations.

###### Data availability statement

The data will be made available at http://www.mrc-cbu.cam.ac.uk/publications/opendata

## 3. Results

### 3.1. Behavioural results

Percentage of responses given revealed a non-significant effect of “Task” [F (1, 44) = 0.163, *p* = .688, *η*p^2^ = .004], indicating that participants gave the same number of responses in the semantic (mean = 79.3 %, SD = 13) and non-semantic tasks (mean = 77.6 %, SD = 13.9).

The RT data revealed a non-significant main effect of “Task” [F (1, 44) = 1.930, *p* = .172, *η*p^2^ = .042] but a significant effect of “Condition” [F (2, 88) = 40.555, *p* < .001, *η*p^2^ = .480]: NC and LC conditions were both slower than HC conditions (*p* = .008 and *p* = .019, respectively), whilst NC and LC conditions did not differ (*p* > .999). A significant interaction between “Condition” and “Task” [F (2, 88) = 20.575, *p* < .001, *η*p^2^ = .319] revealed some RT differences between conditions in the two tasks (see **Figure 2**).

**Figure 2.**
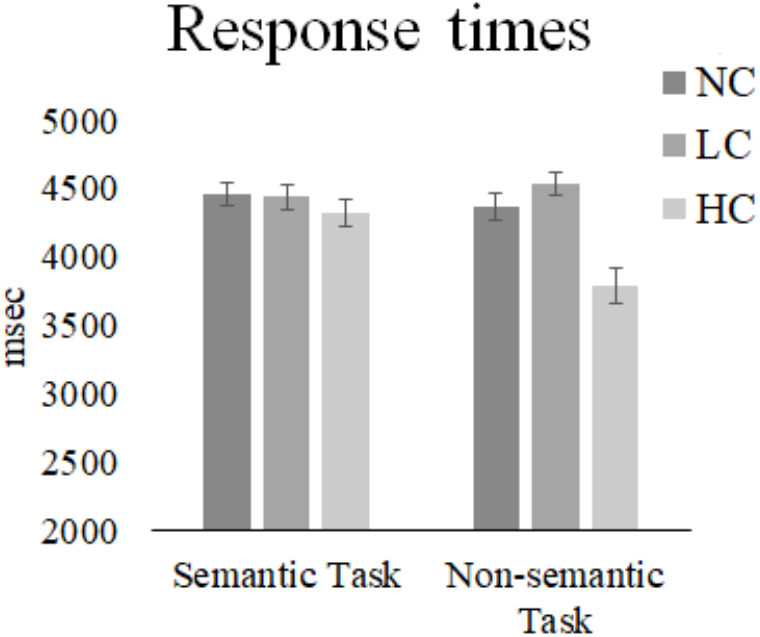
Behavioural results. RTs in semantic and non-semantic tasks for NC, LC and HC target conditions. Error bars correspond to Standard Error (SE).

Importantly and in line with the intended experimental design, a significant main effect of “Condition” was observed in both tasks. An ANOVA applied to data from the semantic task, revealed that conditions differed [F (2, 42) = 5.491, *p* = .008, *η*p^2^ = .207]: the HC condition was faster than both LC and NC conditions (*p* = .05 and *p* = .02, respectively), whilst LC and NC did not differ (*p* > .999). A second ANOVA, including data only from the non-semantic task, also revealed a main effect of “Condition” [F (2, 46) = 38.561,*p* < .001, *η*p^2^ = .626]: NC and LC conditions elicited slower responses than HC conditions (both *ps* < .001), whilst LC and NC did not differ (*p* = .101).

### 3.2. fMRI results

#### 3.2.1. GLM results

##### 3.2.1.1. Whole brain results

###### Context integration in semantic and non-semantic tasks

We first identified which brain regions were engaged during contextual integration, irrespective of task (main effect and conjunction). Then, we examined which brain regions showed an interaction with task.

The main effect of context integration (HC & LC *versus* NC) revealed significant neural activity in the expected fronto-parietal network (see **Figure 3** and **Table S1**), as well as in other brain regions such as the ventro-medial prefrontal cortex, a core area within the Default Mode Network (DMN). Formal conjunction analysis for the same contrast revealed that left inferior parietal cortex, and particularly a cluster including both ventral and dorsal portions of the left AG (peaks of maximal activity corresponding to the MNI coordinates: x = −39, y = −60, z = 36), was similarly involved in semantic and non-semantic context integration (see **Figure 3**).

**Figure 3.**
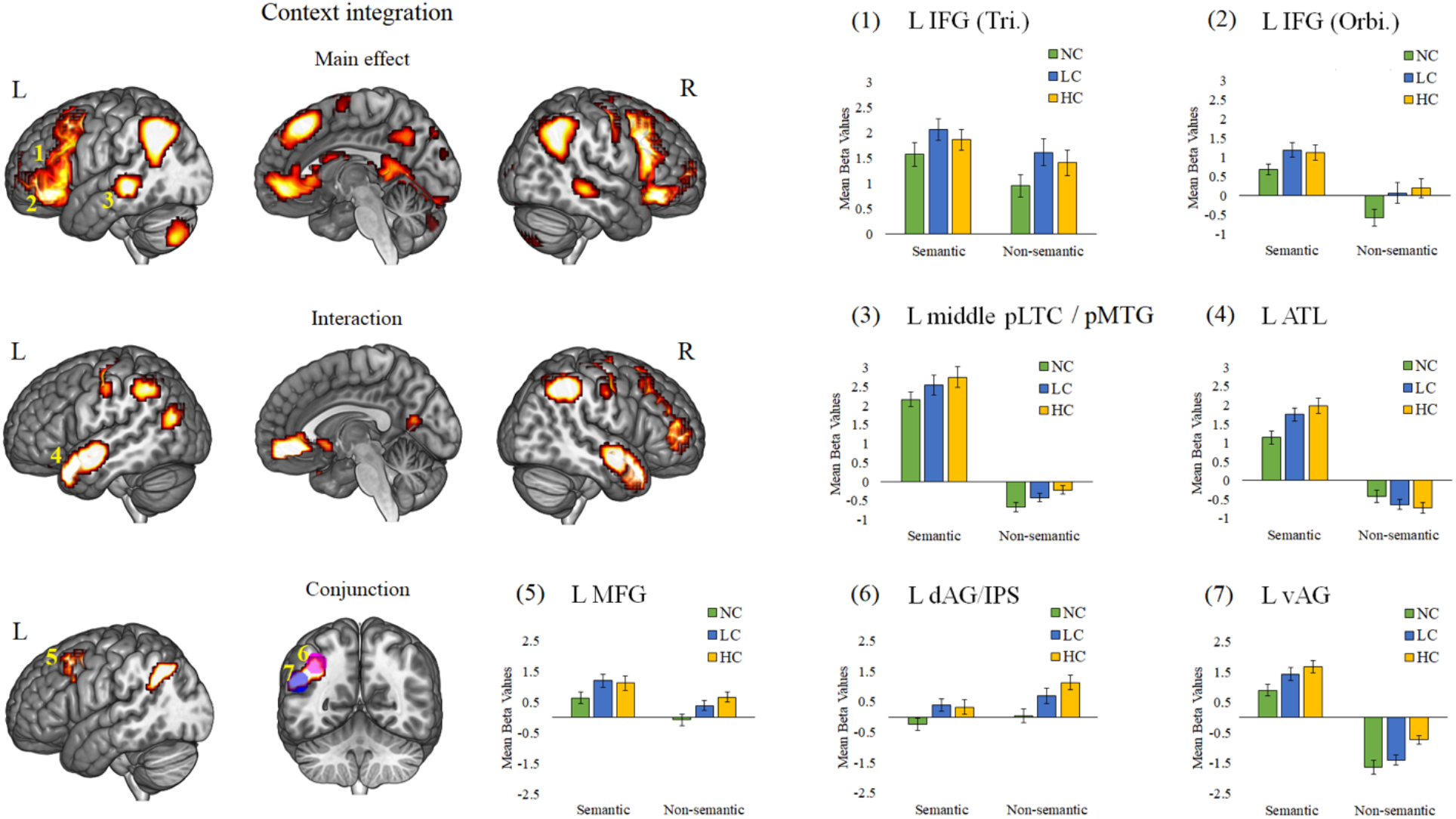
Main effect, interaction, and conjunction analysis results for the context integration effect (HC & LC *versus* NC) in semantic and non-semantic tasks. A voxel-wise level significance threshold was set at *p* < .001 with a FWE correction applied at the critical cluster level at *p* < .05. ROI results reflect mean beta values extracted from frontal, parietal and temporal brain regions. The specific MNI coordinate of each ROI corresponds to cluster activation peaks derived from whole-brain univariate analyses (see **Table 1**). Error bars correspond to SE. *Abbreviations: L = left; R = right; LC = low congruent; HC = high congruent; NC = no-context; IFG = Inferior Frontal Gyrus; Tri = Pars Triangularis; Orbi. = Pars Orbitalis; pLTC/pMTG = posterior lateral temporal cortex/posterior middle temporal gyrus; ATL = Anterior Temporal Lobe; MFG = Middle Frontal Gyrus; dAG/IPS = dorsal Angular Gyrus/Intra-Parietal Sulcus; vAG = ventral angular gyrus*.

###### Context update in semantic and non-semantic tasks

Then we explored which brain regions are important when it is hard to generate contextually integrated representations (LC) in general (i.e., irrespective of task) (main effect) and which brain regions, instead, show task conditional effects (interaction).

The main effect of context update or *hard* contextual integration (LC *versus* HC) revealed significant neural activity in superior parietal cortex and left ventral AG and dorsal AG/IPS (see **Figure 4** and **Table S1**). The results also revealed significant neural responses in precentral gyrus, right insula (anterior and posterior), as well as right putamen (see **Figure 4** and **Table S1**). These regions, however, were not recruited to the very same extent in the two tasks, as indicated by non-significant conjunction analysis results.

**Figure 4.**
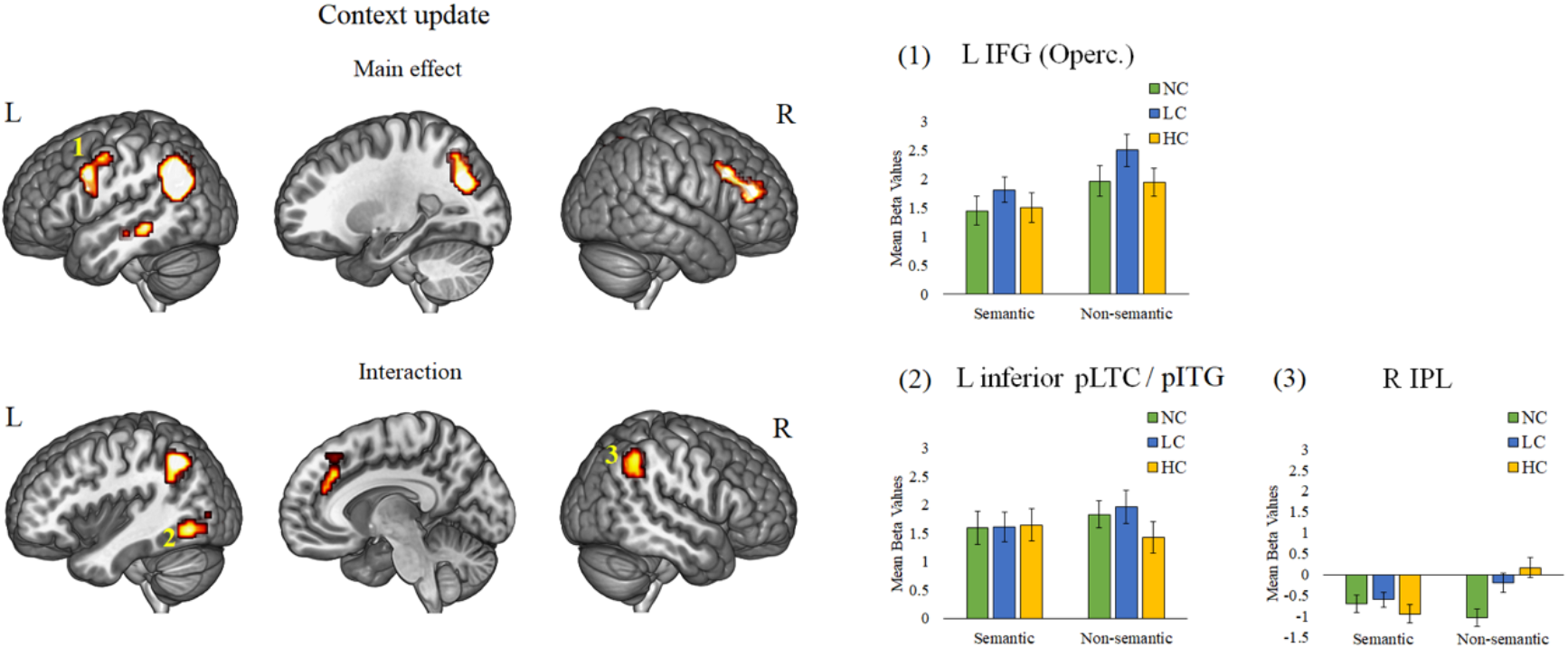
Main effect and interaction for the context update effect (LC *versus* HC) in semantic and non-semantic tasks. A voxel-wise level significance threshold was set at *p* < .001 with a FWE correction applied at the critical cluster level at *p* < .05. ROI results reflect mean beta values from left inferior frontal gyrus (Pars Opercularis), right inferior parietal lobe (IPL) and the left inferior posterior lateral temporal cortex/posterior inferior temporal gyrus (L inferior pLTC/pITG), derived from whole-brain univariate analyses (see **Table 1**). Error bars correspond to SE. *Abbreviations: L = left; R = right; LC = low congruent; HC = high congruent; IFG = Inferior Frontal Gyrus; Operc. = Pars Opercularis*.

Interestingly, the interaction results revealed significant neural activity in bilateral inferior parietal lobes, frontal superior medial lobe and left inferior posterior occipito-temporal cortex (see **Figure 4** and **Table S1**).

##### 3.2.1.2. ROI results

With this analysis, we explored the task-engagement profile of key frontal, parietal and temporal brain regions revealed by whole-brain results. We computed ROI analysis using spheres of 10mm radius centered around the peak of activation within the clusters revealed by the GLM analysis.

Discrete regions within left IFG, left pLTC and left AG, might show functional dissociations depending on the type of stimuli and task difficulty (see Introduction). Accordingly, we selected different frontal (left IFG Pars Opercularis, Triangularis and Orbitalis), temporal (left middle pLTC/pMTG and left inferior pLTC/pITG) and parietal (anterior ventral AG and dorsal AG/IPS) ROIs for these analyses (see **Table 1**), and then we compared the patterns of activation across semantic and non-semantic tasks.

**Table 1.**
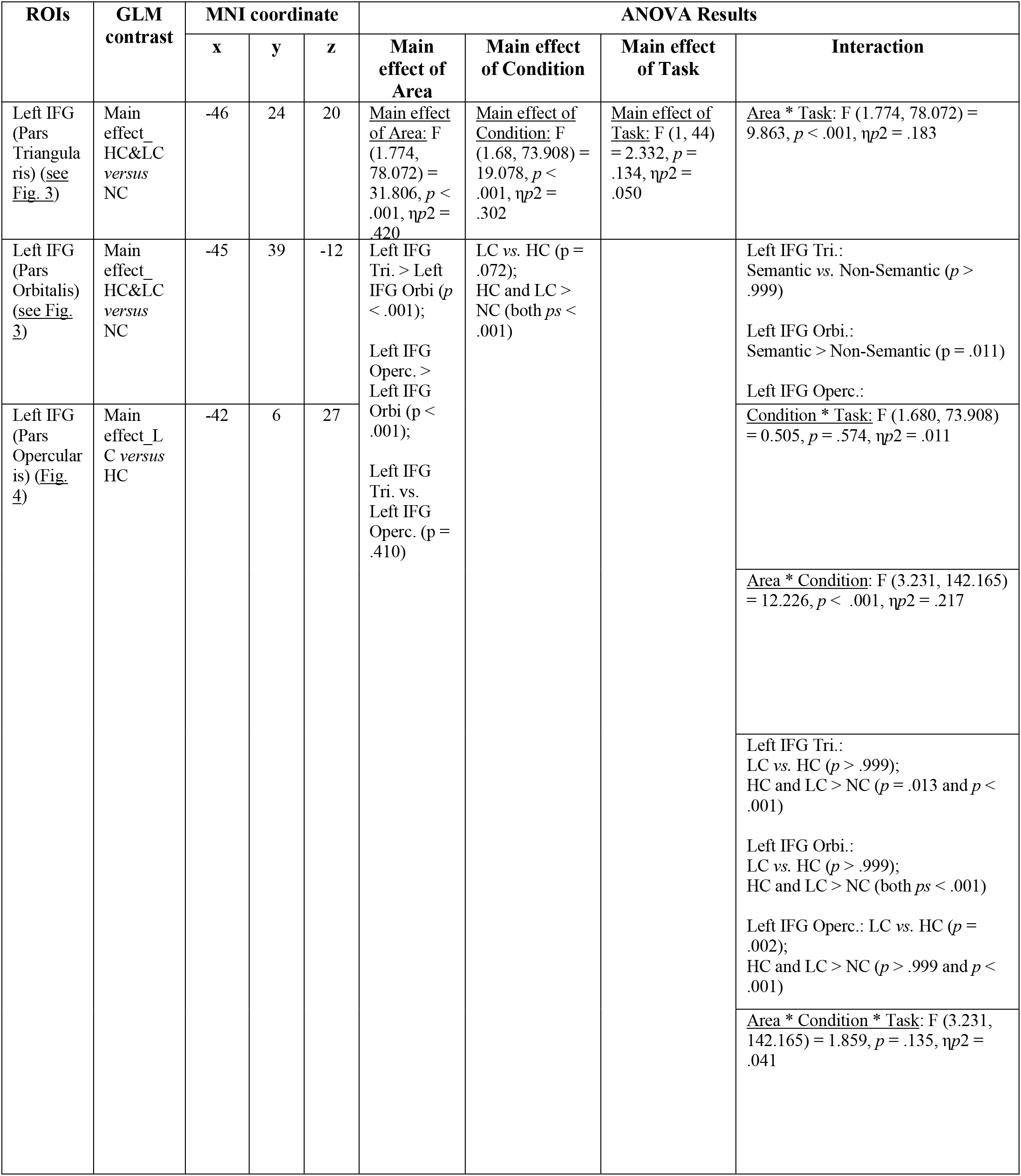
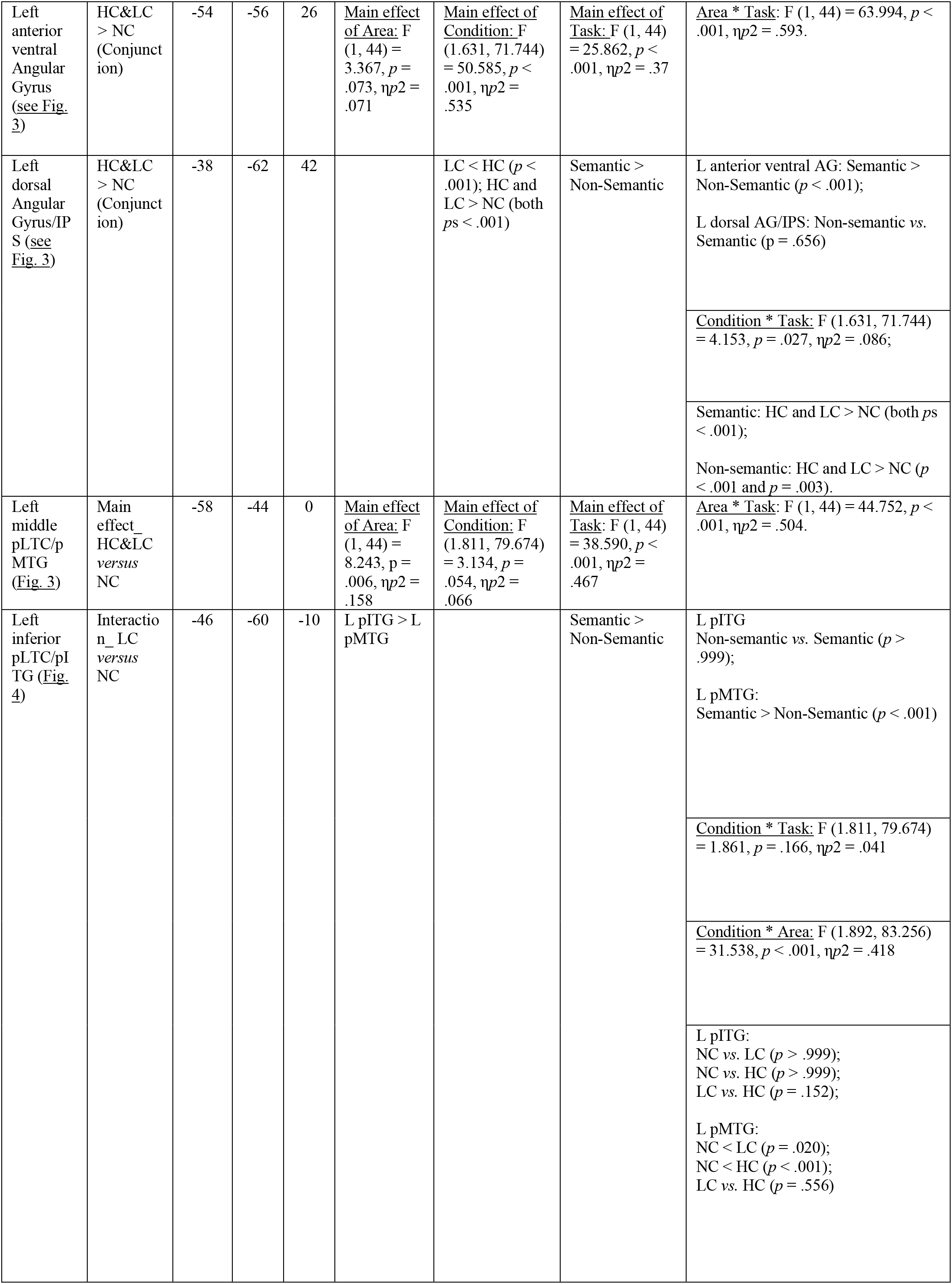
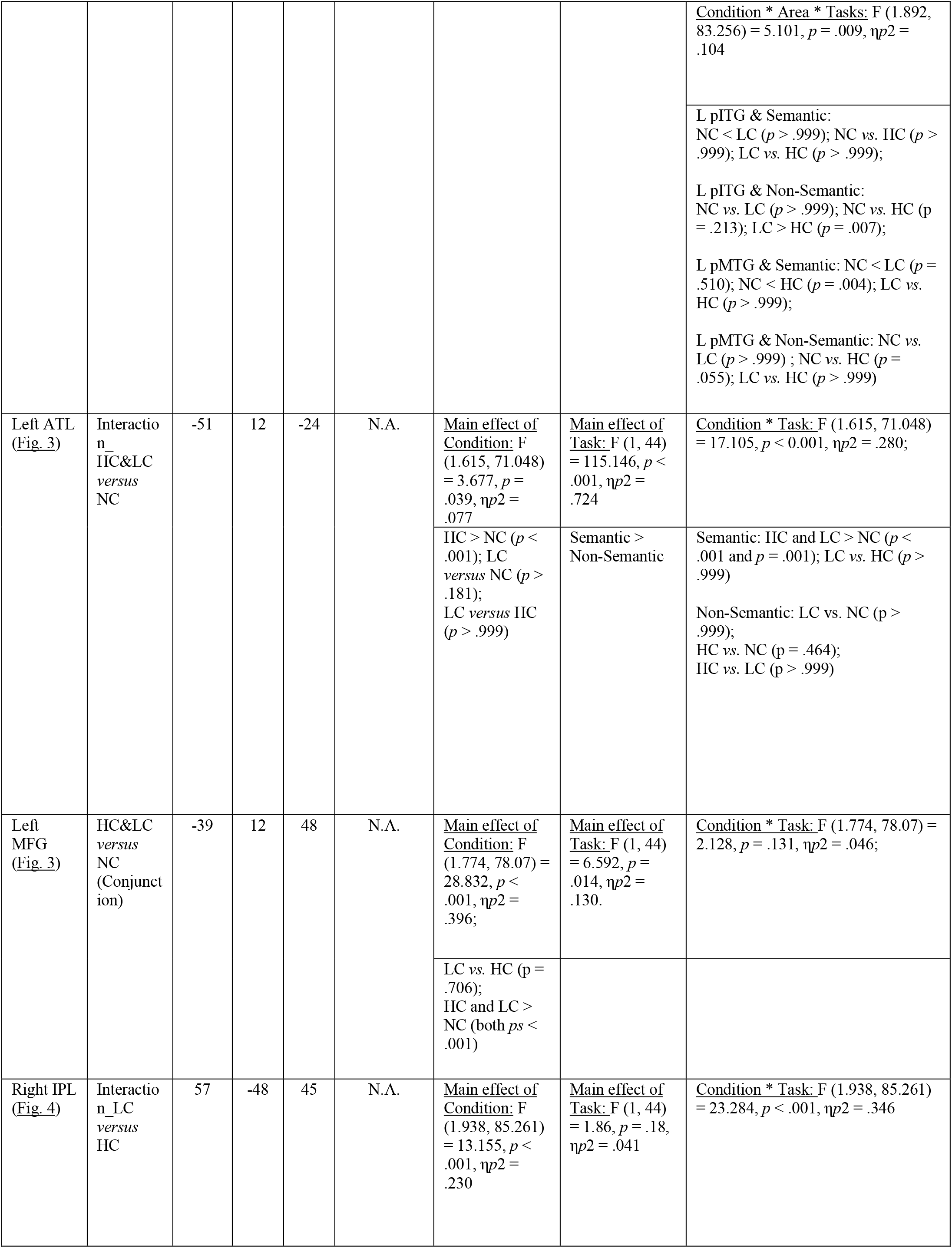
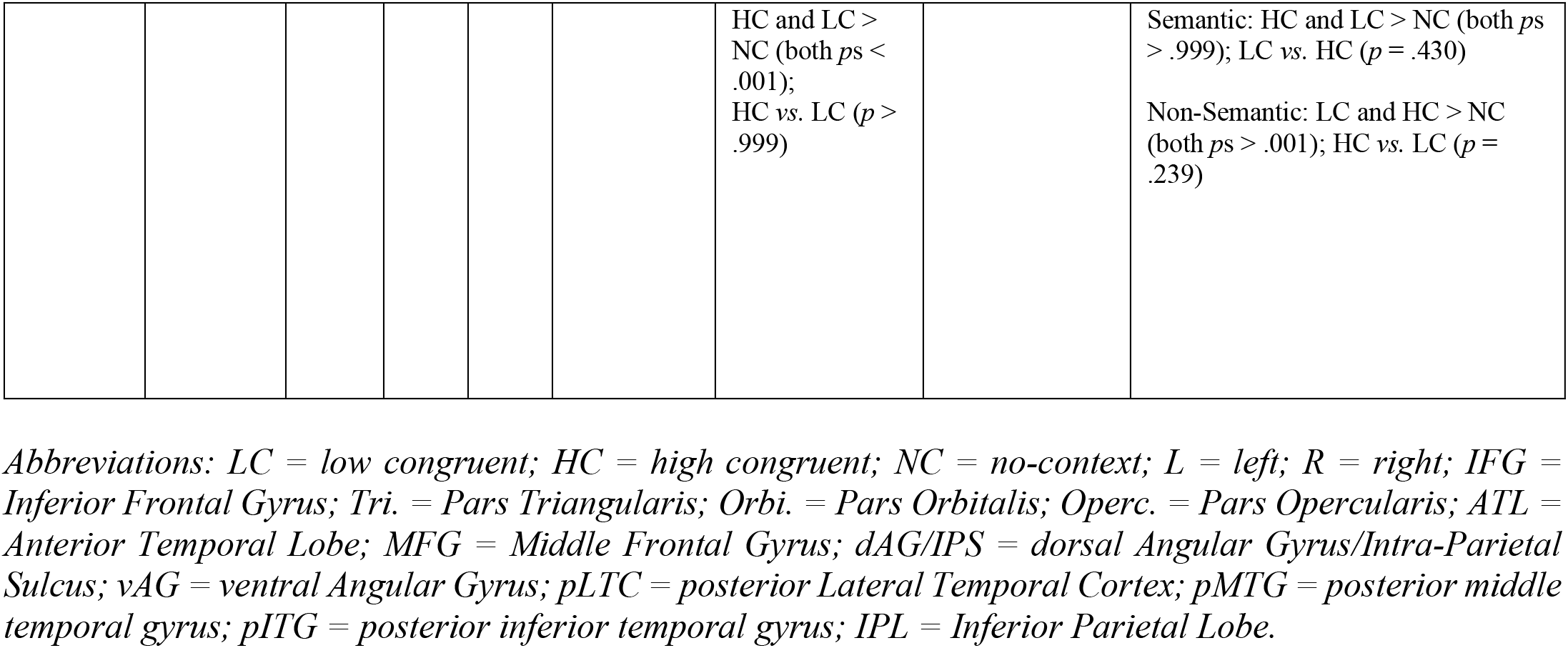
Results from the ROI analysis. Bonferroni correction for multiple comparisons was applied to all post-hoc comparisons. Furthermore, correction for non-sphericity (Greenhouse–Geisser procedure) was applied to the degrees of freedom and *p* values associated with factors having more than two levels.

Finally, we also computed ROI analysis on middle frontal gyrus (MFG) and ATL revealed by whole-brain results (see **Table 1**). Note that in this case our goal was not to compare these brain regions to each other, but just to examine the pattern of activations within each ROI for semantic *versus* non-semantic tasks. Thus, we ran two separate ANOVAs.

The ROI results are reported in **Table 1**. To summarise the most important results, we found that different left IFG subregions were engaged to different extents in semantic and non-semantic tasks. That is, left IFG Pars Triangularis and Pars Opercularis were similarly recruited across tasks, for context integration and context update respectively. Instead, left IFG Pars Orbitalis showed domain-specific responses: A positive pattern of engagement was found in the semantic task, but not in the non-semantic task. In this latter, left IFG Pars Orbitalis’s neural responses were only negative (NC) or negligible (LC and HC), as revealed by one sample *t*-tests (NC: *t* (23) = −2.663, *p* = .014; LC: *t* (23) = 0.257, *p* = .8; HC: *t* (23) = 0.762, *p* = .454) (see **Figure 3**).

A similar dissociation was found in left AG (see **Figure 3**). Left anterior ventral AG was more engaged in semantic *versus* non-semantic task conditions (see **Table 1**), whilst left dorsal AG/IPS was similarly recruited across both tasks. Note that, however, left dorsal AG/IPS was positively engaged only during LC and HC conditions (*t* (23) = 2.66, *p* = .014 and *t* (23) = 2.66, *p* < .001, respectively), confirming the importance of this region for buffering contextual information. Finally, we found that MFG and ATL were both more engaged in semantic *versus* non-semantic task conditions (see **Table 1**).

#### 3.2.2. Task group spatial ICA results

##### 3.2.2.1. Task-related functional networks

ICA identified 26 independent components, of which 5 exhibited significant sensitivity to the task conditions in both semantic and non-semantic tasks: these were a semantic/language network (SLN), an anterior salience network (ASN), left and right executive control networks (LECN and RECN) both including front-oparietal regions. Finally, ICA identified also a DMN component (**see Figure 5**).

**Figure 5.**
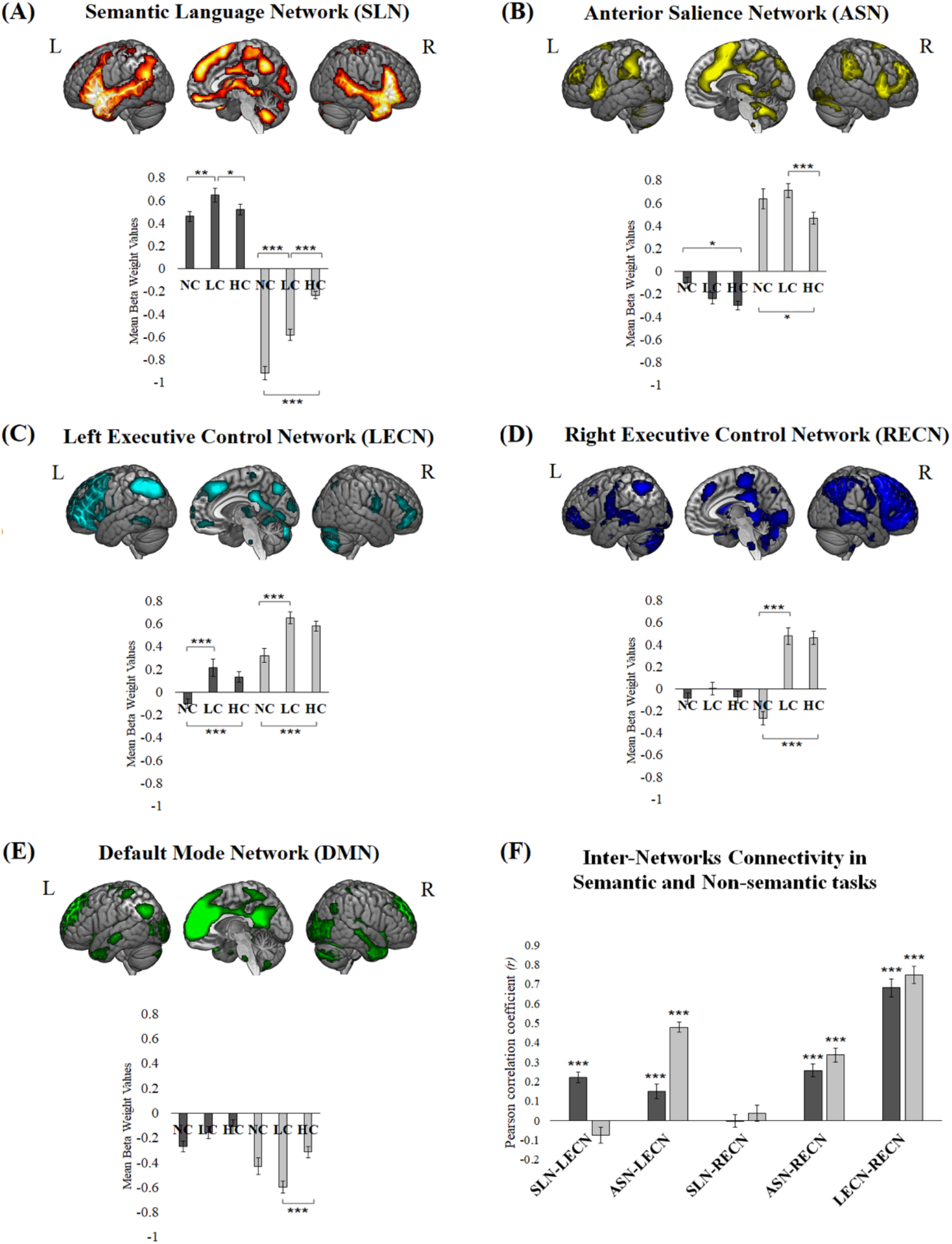
Task-related functional networks. (A) Semantic Language Network (SLN); (B) Anterior Salience Network; (C) left Executive Control Network (LECN); (D) right Executive Control Network (RECN); (E) Default-Mode Network (DMN). ROI results reflect mean beta weight values for each condition (NC, LC and HC), where dark grey bars and light grey bars refer to semantic and non-semantic tasks, respectively. (F) Inter-Networks Connectivity for each task. Asterisks positioned over the bars indicate significant inter-network correlations. Error bars correspond SE. *Abbreviations: L = left; R = right; LC = low congruent; HC = high congruent; NC = no-context*.

A first observation is that the spatial maps of LECN and RECN include similar brain regions as those identified with the context integration univariate results. We conducted a repeated-measures ANOVA on the beta weight values of each component to examine context integration and context update across tasks. The results revealed a significant interaction between “Network” (SLN, ASN, LECN, RECN, and DMN), “Condition” (NC, LC, and HC), and “Task” (semantic and non-semantic), which suggests a different engagement of the networks in the different conditions and tasks [F (8,352) = 18.337,*p* < .001, *η*p^2^ = .294].

The SLN was differently engaged in the two tasks (see **Figure 5**). In the semantic task, SLN showed increased responses for LC conditions as compared to both HC and NC conditions (*p* = .035 and *p* = .005, respectively), whilst the difference between HC and NC was not significant (*p* = .895). In the non-semantic task, there were decreased, rather than increased, responses in SLN as compared to rest baseline. Thus, HC conditions showed less deactivation as compared to LC and NC conditions (both *ps* < .001), whilst the LC conditions showed less deactivation than the NC conditions (*p* < .001).

The ASN showed an opposite pattern of results to the SLN. Neural responses were positive in the non-semantic task, but negative in the semantic task. In the semantic task, NC conditions differed from HC conditions (*p* = .013). There were no significant differences between LC and HC conditions (*p* = .436), nor between LC and NC conditions (*p* = .124). In the non-semantic task, NC and LC conditions showed more positive responses as compared to HC conditions (*p* = .030 and *p* < .001), whilst the LC and NC conditions did not differ (*p* = .757).

The LECN was similarly engaged in the two tasks. In the semantic task, it showed increased positive responses when the semantic context was available for integration (LC > NC and HC > NC, both *ps* < .001), but no difference between LC and HC was observed (*p* = .471). Also in the non-semantic task, increased positive responses were observed when context integration was possible (LC > NC and HC > NC, both *ps* < .001). As above, no difference between LC and HC was observed (*p* = .515). The RECN showed a similar context integration effect as for LECN, but only in the non-semantic task (HC > NC and LC > NC, both *ps* < .001; HC *versus* LC, *p >* .999).

Finally, unlike the other components, the DMN was deactivated with respect to rest. Conditions differed only in the non-semantic task and reflected differential de-activations (non-semantic task: NC *versus* LC: *p* = .054; NC *versus* HC: *p* = .318; HC > LC: *p* < .001).

##### 3.2.2.2. FNC analysis

Given our interest in exploring functional interactions between the domain-specific (SLN, ASN, RECN) and domain-general networks (LECN), we computed a FNC analysis (see **Figure 5F**). An intriguing pattern of results emerged. The strength of correlations between domain-specific and domain-general networks changed as a function of the task, as revealed by a repeated-measures ANOVA. In detail, in this analysis “Task” was a between-subjects factor with two levels (semantic and non-semantic) and “Type of FNC” a within-subjects factor with five levels (SLN-LECN, SLN-RECN, ASN-LECN, ASN-RECN and LECN-RECN). Bonferroni correction for multiple comparisons was applied to assess statistically significant effects. Correction for non-sphericity (Greenhouse–Geisser procedure) was applied to the degrees of freedom and *p* values associated with factors having more than two levels.

The results revealed a significant interaction between “Task” and “Type of FNC” [F (2.566, 112.886) = 19.822, *p* < .001, *η*p^2^ = .311], indicating that the significant positive correlation observed between time courses of LECN and SLN was stronger in the semantic *versus* non-semantic tasks (*p* < .001). Instead, the opposite pattern was observed for LECN-ASN coupling, i.e., stronger positive correlation in the non-semantic task as compared to the semantic (*p* < .001). Interestingly, SLN-RECN, ASN-RECN and LECN-RECN did not show any difference between tasks (*p* > .999) (see **Figure 5F**).

#### 3.2.3. Summary of the results

The main effect of context integration revealed strong brain activations within the semantic control network (Lambon Ralph *et al*. 2017; Jackson 2020), including left IFG, left pLTC, as well as the left AG. A closer look to these results revealed dissociations in the patterns of engagement within these regions. For instance, left IFG (Pars Orbitalis), left anterior ventral AG and left middle pLTC/pMTG were uniquely engaged during integration of semantic information. Instead, IFG Pars Triangularis and left dorsal AG/IPS did not show these domain-specific effects, but rather a domain-general role for generating context-integrated representations.

Interestingly, the main effect of context update/hard contextual integration revealed significant activations in different portions of the frontal and parietal lobes, as compared to those revealed by the main effect of context integration. In fact, left posterior IFG (Pars Opercularis) as well as the superior parietal cortex were involved in updating the contextual representation, irrespective of the type of stimuli (**Figure 4** and **Table S1**). Interestingly, the inferior pLTC/pITG showed a context update effect but only for non-semantic stimuli. Finally, the GLM and ROI results also revealed that left superior lateral ATL and left pMTG were uniquely involved in semantic contextual integration (see **Figure 3** and **Table 1**).

ICA results showed that whilst LECN supports domain-general context integration, the update of contextual information is supported by domain-specific networks (ASN and SLN). Interestingly, besides LECN, also a RECN is involved in context integration, but only for the generation of non-semantic contextual representations. Finally, interactions between LECN and domain-specific networks are modulated by the type of task, and therefore, the nature of the stimuli involved during context integration.

## 4. Discussion

The present fMRI study investigated the extent to which neural processes reflecting context integration and context update are domain-specific *versus* domain-general. Further exploration of whole-brain results via ROI analysis focused particularly on the role of different subregions of left IFG, left pLTC and left AG. In fact, these regions are functionally heterogenous, with ongoing debates regarding (1) which sub-regions are implicated in semantic processing and beyond (Seghier et al. 2010; Seghier 2013; Diachek *et al*. 2020; Hodgson *et al*. 2021; Humphreys *et al*. 2021; Wehbe et al. 2021) and (2) the extent to which their involvement in semantic tasks might reflect methodological artifacts (Fedorenko and Shain 2021).

We hypothesized to find both shared (left IFG Pars Opercularis and Triangularis, left inferior pLTC/pITG, and left dorsal AG/IPS) and distinct (left IFG Pars Orbitalis, middle pLTC/pMTG and anterior ventral AG for semantic stimuli only) regions involved in context integration and update. Aligning with our predictions, we found that left IFG, left pLTC and left AG functionally fractionate, as they revealed both domain-specific and domain-general subregions for context integration and update. The results, which are discussed below, provide the first characterization of the neural processes during naturalistic semantic processing along the domain-specificity dimension, and at the same time, bring new insights on the role of left inferior pLTC/pITG and left dorsal AG/IPS for semantic cognition.

### Left IFG

The left IFG Pars Opercularis was sensitive to control demands during context integration (LC > HC), and this effect was independent from the type of task/stimulus. This is consistent with previous studies that have found left IFG Pars Opercularis to be more active when sentence meaning was ambiguous or implausible as compared to plausible (Rodd et al. 2005; Willems et al. 2009; Desai et al. 2010; Obleser and Kotz 2010). This neural effect has been interpreted in many different ways (Price 2012). Some have proposed that this could reflect processing violations of the “what” related predictions and integration/update of input (Obleser and Kotz 2010). Others have related this effect to increased verbal working memory demands (Koelsch et al. 2009). Our results do not seem to favour the latter hypothesis. In fact, we did not observe any difference between conditions that require (HC) *versus* those that do not require (NC) to maintain information in working memory to allow context integration. Thus, it is possible that the context update effect observed in left IFG Pars Opercularis reflects the consequences of a detection of violation of prediction. Our data also show that left IFG Pars Opercularis was similarly engaged in semantic and non-semantic tasks. This is in accord with studies that revealed incongruency effects in this sub-region, not only in language comprehension, but also in music and action perception (Koelsch et al. 2002; Tillmann et al. 2006; Bianco et al. 2016; Siman-Tov et al. 2019).

Since neural responses observed in left Pars Opercularis were quite specific for the LC condition (i.e., did not track task difficulty) and, at the same time, not restricted to the verbal/semantic domain, it is likely that this subregion supports domain-general processes for context integration, that might come into play only under increased task demands, e.g., during the update or reset of an evolving representation.

Also left IFG Pars Triangularis was recruited by semantic and non-semantic tasks alike, a result that aligns with our hypothesis that this brain region would reflect domain-general processes for context integration. Indeed, previous studies have found that this subregion is not only implicated in the control of semantic information (Badre and Wagner 2002; Gold et al. 2005; Gold et al. 2006). For instance, competition in episodic and working memory tasks also increases activation of this brain region (Badre and Wagner 2002; Dobbins and Wagner 2005).

Somewhat unexpectedly, the left IFG Pars Triangularis did not show enhanced neural responses during context update or LC conditions. Incongruency effects in left IFG Pars Triangularis have been mainly observed in tasks where retrieval of information occurred with little or no contextual support (Rodd *et al*. 2005; Whitney, Jefferies, et al. 2011; Price 2012; Humphreys *et al*. 2019; Hodgson *et al*. 2021). Therefore, it is possible that those incongruency effects reflected task-specific demands rather than semantic control processes typically required in context integration itself.

As expected, left IFG Pars Orbitalis exhibited domain (semantic)-specific responses and was sensitive to context integration (HC & LC > NC). This observation is in accordance with a recent meta-analysis study where left IFG Pars Orbitalis - unlike Pars Opercularis and Triangularis - was modulated by semantic control demands but not by other type of non-semantic demands. Accordingly, this anterior and ventral subregion of the left IFG lies outside the MDN (Duncan 2010; Fedorenko *et al*. 2013; Assem *et al*. 2020; Duncan *et al*. 2020). Thus, one possibility is that left IFG Pars Orbitalis is specialized for the control and manipulation of semantic information (Wagner et al. 2001; Devlin et al. 2003; Badre et al. 2005; Gough et al. 2005). In contrast, more posterior-dorsal subregions (Pars Triangularis and Pars Opercularis) seem to support domain-general computations. Alternatively, left IFG’s activity would reflect the same neuro-computation (e.g., context integration) and the graded differences might depend on variations in functional and structural connectivity with areas and/or networks that support domain-specific processes. For instance, left IFG Pars Orbitalis is directly connected via uncinate fasciculus to the ATL (Binney et al. 2012), a neural hub for semantic representations (Patterson et al. 2007; Lambon Ralph *et al*. 2017). Instead, left IFG Pars Opercularis and Triangularis are directly connected to MDN brain regions, such as the dorsolateral prefrontal cortex (Jung et al. 2022), which are activated across different domains when task demands increase. In our data, left IFG activations during context integration and update overlapped substantially across tasks. This was found not only in the whole brain results (ANOVA), but also in the functional networks revealed by ICA, where domain-specific (SLN and ASN) and domain-general networks (LECN) overlapped in left IFG, a result that does not align neatly with a semantic *versus* domain-general distinction for the functional role of left IFG.

### Left pLTC

Whole-brain analyses revealed that left middle pLTC/pMTG responded similarly to context integration in semantic and non-semantic tasks, yet it was positively engaged in the semantic task only. Accordingly, left middle pLTC/pMTG was embedded in the semantic-specific network (SLN), but not in the network showing domain-general effects (e.g., LECN). These results are in accordance with our hypothesis that left middle pLTC/pMTG would be specialized for the integration of semantic information (Hodgson *et al*. 2021). Like left IFG Pars Orbitalis, left middle pLTC/pMTG did not show increased neural responses when integration demands increased (LC condition). It is possible that this effect was not observed because of differences between our and previous studies (e.g., semantic judgment task, thematic association task) (Whitney, Kirk, *et al*. 2011). In our semantic task, participants were provided with rich semantic contexts, which is very different from previous paradigms where little contextual information was provided during semantic retrieval, potentially increasing the control demands by necessitating additional context-relevant information retrieval.

Interestingly, whole-brain results revealed a completely different pattern of engagement in left inferior pLTC/pITG. This subregion was engaged to the same extent in semantic and non-semantic tasks. However, our results revealed a role for this subregion in context update during the processing of numerical, but not semantic, stimuli. This finding, which is at odds with our initial hypothesis, i.e., to find this subregion similarly engaged in semantic and non-semantic tasks for context update (Hodgson *et al*. 2021), suggests that left inferior pLTC/pITG may not be recruited for semantic control, but only as part of MDN when the task loads strongly on executive control (Fedorenko and Shain 2021). This proposal is also consistent with the fact that left inferior pLTC/pITG was functionally embedded in the LECN but not the SLN.

### Left AG

Whole-brain results revealed a functional overlap across tasks in the left AG, and especially left dorsal AG/IPS, implicating this region in context integration processes beyond the semantic domain. Accordingly, left dorsal AG/IPS was recruited within a fronto-parietal network (LECN), which was sensitive to context integration in both tasks.

We also revealed some domain-specific responses within the left AG. As expected, left anterior ventral AG was positively engaged (as compared to rest) only when context integration involved semantic stimuli. Note that the dissociation observed between neural responses in left dorsal AG/IPS and left anterior ventral AG cannot be related to differences in task difficulty, as there was no RT difference between the two tasks. Furthermore, both subregions showed context integration effects that did not seem to track difficulty-related differences between conditions.

The domain-specific and domain-general (conjunction results) neural results presented above are consistent with the proposal that the core function of left AG is a functionally-graded domain-general buffer, with subregions showing different patterns of engagement depending on their pattern of connectivity to various domain-specific systems (Humphreys and Lambon Ralph 2015; Humphreys et al. 2022). Accordingly, left anterior ventral AG which is functionally and anatomically connected to the language network (see SLN) (Makris et al. 2013; Humphreys *et al*. 2022), showed strong positive responses during the integration of verbal, but not numerical information.

We also tested the hypothesis of a differential involvement of dorsal and anterior ventral AG regions depending on the context integration demands. Specifically, we hypothesized that an update of context (LC > HC) would require top-down control processes, implemented by left dorsal AG/IPS (Corbetta and Shulman 2002; Noonan, Jefferies, Visser and Lambon Ralph 2013; Humphreys and Lambon Ralph 2015; Humphreys and Lambon Ralph 2017). Conversely, we predicted stronger neural responses for HC *versus* LC conditions in left ventral AG, reflecting automatic bottom-up buffering processes (Humphreys and Lambon Ralph 2015). Interestingly, we observed stronger responses for HC as compared to LC conditions in both subregions (see **Figure 3**). One possibility is that left dorsal AG/IPS’s sensitivity to task demands reported by previous studies did not primarily reflect buffering-related neural processes, but rather top-down attentional control processes imposed by a secondary task (e.g., a memory probe, comprehension, or sentence judgment task).

Overall, the findings reviewed above are in accordance with previous reports showing a link between left AG’s activity and integration of contextual information in language tasks (van der Linden et al. 2017; Branzi, Humphreys, *et al*. 2020; Branzi, Pobric, *et al*. 2021). The current study goes further by establishing that left dorsal AG/IPS reflects domain-general processes for context integration (Humphreys and Lambon Ralph 2015, 2017; Humphreys *et al*. 2019; Ramanan and Bellana 2019). Our results have implications for the debate on the role of left AG in semantic cognition. In fact, in contrast with the proposal of the domain-general buffer’s role, other researchers have argued that left AG would reflect a supramodal conceptual semantic hub (Binder et al. 2009). Our results are not compatible with this view. In fact, conjunction analysis results revealed that left AG and especially dorsal AG/IPS was recruited to the same extent in both tasks.

### Domain-general and domain-specific functional networks

The ICA results revealed a fronto-parietal network (LECN), including working memory as well as MDN areas (Hodgson *et al*. 2021). The LECN was similarly recruited for semantic and non-semantic context integration. In contrast, a language network (i.e., SLN) including anterior and posterior temporal areas, left anterior ventral AG and left IFG, was positively engaged during the semantic task, and especially for context update. This network was suppressed during the non-semantic task. ICA revealed another domain-specific network, the ASN, primarily composed of the anterior insula and dorsal anterior cingulate cortex (Menon and Uddin 2010; Menon 2011). This network was strongly engaged during the non-semantic task for context update but suppressed during the semantic task. Finally, our results revealed that interactions between LECN and the domain-specific networks (i.e., SLN and ASN) were modulated by the type of task, and therefore, the nature of the stimuli involved in context integration.

The results from the ICA complement the whole-brain results and deepen our understanding of the neural mechanisms behind context integration and update. The LECN, consisting of fronto-parietal brain regions, probably reflects domain-general processes to build contextual representations. We propose that this network is engaged during online buffering and integration of information over time. The LECN also interacts with other two domain-specific networks, the ASN and SNL, depending on the task. It has been suggested that the ASN plays an important role in detection of unexpected or salient stimuli and the subsequent engagement of the LECN for working memory and higher-order cognitive control (Menon 2011). In contrast, the SLN has been associated to language-specific neurocomputations, including tracking of coherence and the detection of semantic violations (Branzi, Humphreys, *et al*. 2020; Diachek *et al*. 2020; Fedorenko and Shain 2021). The SNL’s activity is also accompanied by the activity of fronto-parietal brain regions, especially when task demands increase. Accordingly, it is possible that ASN and SLN detect violations of domain-specific expectations. Then, the need to update the ongoing representation might determine the additional recruitment of the LECN, which would aid integration and update of the ongoing representations within domain-specific areas (e.g., ATL). The observed interactions between domain-specific networks and LECN are in accordance with the hypothesis that domain-specific and domain-general networks work in tandem for context integration.

Finally, ICA also revealed a RECN, which is bilateral yet asymmetric unlike the LECN which is more strongly left lateralized. This network was uniquely engaged in the non-semantic task, in accord with the lack of functional interactions between RECN and SLN. It is possible that the non-semantic task requires processes not shared with the linguistic domain for the integration of numerical information. This might explain the hemispheric fractionation (RECN and LECN) of the executive control network in the non-semantic task only.

### Conclusion

Our results suggest that semantic control and domain-general control are dissociable (IFG, AG, pMTG) and thus that semantic control processes are not fully analogous to other types of cognitive demands (e.g., Gao et al. 2021). Using ICA, we revealed a functional segregation between LECN and SLN during language, in accordance with the idea that MDN brain regions might not reflect core-linguistic operations (e.g., update of the semantic gestalt) (Diachek *et al*. 2020; Fedorenko and Shain 2021; Wehbe *et al*. 2021). Nevertheless, our results indicate that these control systems are highly interactive. This is in line with multiple lines of research. For instance, after stroke, domain-general executive function skills are important predictors of the aphasia recovery (Geranmayeh et al. 2017). Furthermore, stimulating regions from the MDN improves language acquisition (Sliwinska et al. 2017), a finding which is consistent with evidence of transfer effects between language control and domain-general executive control in healthy and patient populations (e.g., Cattaneo et al. 2015; Timmer et al. 2019). Our findings complement this evidence by showing that during naturalistic language processing the SLN and the LECN strongly interact. Future studies are needed to understand the functional relevance of these interactions, and *if* and *when* these interactions become *necessary* for language cognition.

## Acknowledgements

This research was supported by a Medical Research Council Programme Grant (MR/R023883/1), an European Research Council Advanced Grant (GAP: 670428 - BRAIN2MIND_NEUROCOMP), a Postdoctoral Fellowship from the European Union’s Horizon 2020 research and innovation programme, under the Marie Sklodowska-Curie grant agreement No 658341, and Medical Research intramural funding (MC_UU_00005/18. For the purpose of open access, the UKRI-funded authors have applied a Creative Commons Attribution (CC BY) licence to any Author Accepted Manuscript version arising from this submission.

## Supplementary Information

**Table S1.** The locations of the activation peaks (MNI coordinates) from the GLM analyses.

| Contrast | Cluster size | *F* | x | y | z | Peak Location |
| --- | --- | --- | --- | --- | --- | --- |
| Main effect<br>HC&LC *versus* NC | 1145 | 76,82506 | -42 | -60 | 45 | L Angular Gyrus |
|  |  | 58,13088 | -57 | -60 | 27 | L Angular Gyrus |
|  |  | 53,776 | -57 | -30 | -6 | L Middle Temporal Gyrus |
|  | 216 | 54,1929 | 15 | -78 | -33 | R Cerebelum Crus 2 |
|  |  | 27,16881 | 27 | -78 | -45 | R Cerebelum Crus 2 |
|  |  | 23,42683 | 36 | -75 | -48 | R Cerebelum Crus 2 |
|  | 2009 | 53,83453 | 0 | 30 | 51 | L Superior Medial Gyrus |
|  |  | 50,30552 | -42 | 15 | 48 | L Middle Frontal Gyrus |
|  |  | 47,54526 | -45 | 39 | -12 | L IFG p. Orbitalis |
|  | 837 | 52,60666 | 66 | -33 | -3 | R Middle Temporal Gyrus |
|  |  | 48,46903 | 45 | -57 | 45 | R Inferior Parietal Lobule |
|  |  | 44,20713 | 45 | -54 | 33 | R Angular Gyrus |
|  | 1114 | 49,67309 | 45 | 18 | 45 | R Middle Frontal Gyrus |
|  |  | 48,10902 | 33 | 54 | 9 | R Superior Frontal Gyrus |
|  |  | 47,31599 | 57 | 24 | 27 | R IFG p. Triangularis |
|  | 364 | 48,06685 | -12 | -78 | -33 | L Cerebelum Crus 2 |
|  |  | 28,65035 | -33 | -75 | -48 | L Cerebelum Crus 2 |
|  | 1036 | 36,18142 | -12 | 12 | 24 | L Caudate |
|  |  | 34,18916 | 3 | 48 | -6 | R Mid Orbital Gyrus |
|  |  | 33,07639 | -18 | 27 | 6 | L Caudate |
|  | 275 | 34,3563 | 48 | -6 | 54 | R Middle Frontal Gyrus |
|  |  | 22,5165 | 51 | -6 | 39 | R Precentral Gyrus |
|  |  | 18,31665 | 3 | 0 | 66 | R Posterior-Medial Frontal |
|  | 178 | 33,08943 | 9 | -57 | 36 | R Precuneus |
|  |  | 18,20664 | -9 | -48 | 36 | L MCC |
|  | 392 | 28,09912 | -3 | -42 | 3 | Lobule I IV Hem |
|  |  | 20,21798 | 6 | -45 | 15 | R Precuneus |
|  |  | 17,6854 | 0 | -87 | -18 | Lobule VI Verm |
|  | 278 | 22,53702 | 18 | -87 | 0 | R Calcarine Gyrus |
|  |  | 17,87527 | 24 | -96 | 9 | R Middle Occipital Gyrus |
|  |  | 16,39382 | 3 | -96 | -3 | L Calcarine Gyrus |
| Interaction<br>HC&LC *versus* NC | 339 | 58,62682 | 48 | 15 | -30 | R Medial Temporal Pole |
|  |  | 51,43274 | 57 | 0 | -18 | R Middle Temporal Gyrus |
|  |  | 49,13137 | 54 | 9 | -24 | R Medial Temporal Pole |
|  | 882 | 53,34651 | 3 | 42 | -15 | R Rectal Gyrus |
|  |  | 37,57184 | -18 | 33 | -15 | L Middle Orbital Gyrus |
|  |  | 23,2614 | -21 | -3 | -24 | L Amygdala |
|  | 398 | 49,75475 | -57 | 9 | -18 | L Temporal Pole |
|  |  | 46,19533 | -51 | 12 | -24 | L Temporal Pole |
|  |  | 45,93649 | -60 | -6 | -9 | L Middle Temporal Gyrus |
|  | 225 | 35,73101 | -42 | -72 | 24 | L Middle Occipital Gyrus |
|  | 314 | 33,91481 | 51 | -48 | 48 | R Inferior Parietal Lobule |
|  | 421 | 33,45044 | 39 | 48 | 0 | R Middle Frontal Gyrus |
|  |  | 21,38867 | 42 | 21 | 48 | R Middle Frontal Gyrus |
|  |  | 19,24164 | 45 | 39 | 30 | R Middle Frontal Gyrus |
|  | 220 | 29,23179 | -45 | -15 | 51 | L Postcentral Gyrus |
|  |  | 16,25564 | -33 | -24 | 66 | L Precentral Gyrus |
|  | 156 | 25,82463 | -51 | -54 | 45 | L Inferior Parietal Lobule |
|  | 149 | 22,18096 | -21 | -63 | 12 | L Calcarine Gyrus |
|  |  | 20,04431 | -9 | -63 | 12 | L Calcarine Gyrus |
|  | 228 | 21,21912 | 39 | -27 | 66 | R Precentral Gyrus |
|  |  | 20,9011 | 51 | -12 | 51 | R Precentral Gyrus |
|  |  | 18,12021 | 27 | -12 | 69 | R Superior Frontal Gyrus |
| Conjunction<br>(Semantic & Non-semantic)<br>HC&LC *versus* NC | 168 | 28,82601 | -39 | -60 | 36 | L Angular Gyrus |
|  |  | 27,06504 | -57 | -57 | 27 | L Angular Gyrus |
|  |  | 25,55533 | -48 | -57 | 30 | L Angular Gyrus |
|  | 143 | 24,34014 | -39 | 12 | 48 | L Middle Frontal Gyrus |
|  |  | 16,59037 | -33 | 18 | 48 | L Middle Frontal Gyrus |
|  |  | 13,19632 | -48 | 15 | 33 | L IFG p. Opercularis |
| Main effect<br>LC *versus* HC | 445 | 56,58667 | -45 | -57 | 27 | L Angular Gyrus |
|  | 208 | 48,38602 | -42 | 6 | 27 | L IFG p. Opercularis |
|  |  | 17,66829 | -48 | -6 | 45 | L Precentral Gyrus |
|  | 540 | 43,77959 | 27 | -69 | 33 | R Superior Occipital Gyrus |
|  |  | 23,0356 | 36 | -42 | 39 | Area hIP1 IPS |
|  |  | 13,88197 | 18 | -54 | 21 | R Precuneus |
|  | 328 | 36,0464 | 51 | 39 | 21 | R Middle Frontal Gyrus |
|  |  | 31,89053 | 30 | 27 | 6 | R Insula Lobe |
|  |  | 26,8577 | 42 | 6 | 30 | R Precentral Gyrus |
|  | 137 | 26,48048 | -21 | -69 | 42 | L Superior Parietal Lobule |
|  |  | 26,02419 | -27 | -69 | 30 | L Middle Occipital Gyrus |
|  | 123 | 24,24963 | -57 | -33 | -9 | L Middle Temporal Gyrus |
|  |  | 21,67479 | -60 | -21 | -9 | L Middle Temporal Gyrus |
|  | 137 | 22,18175 | 39 | 0 | 15 | R Insula Lobe |
|  |  | 18,46788 | 48 | 0 | 12 | R Rolandic Operculum |
|  |  | 13,9542 | 48 | -6 | 21 | R Rolandic Operculum |
| Interaction<br>LC *versus* HC | 206 | 26,86099 | -45 | -57 | 45 | L Inferior Parietal Lobule |
|  |  | 22,4724 | -51 | -54 | 30 | L Angular Gyrus |
|  | 126 | 19,92358 | 57 | -48 | 45 | R Inferior Parietal Lobule |
|  |  | 15,57321 | 45 | -51 | 42 | R Inferior Parietal Lobule |
|  | 142 | 19,23207 | 6 | 30 | 39 | R Superior Medial Gyrus |
|  |  | 16,88592 | -6 | 33 | 33 | L Superior Medial Gyrus |
|  |  | 16,67199 | 0 | 30 | 51 | L Superior Medial Gyrus |
|  | 127 | 18,91905 | -42 | -69 | -9 | L Inferior Occipital Gyrus |
|  |  | 17,12739 | -39 | -84 | -3 | L Middle Occipital Gyrus |

*Abbreviations:* L = left; R= right; p.=Pars; MCC = Middle Cingulate Cortex; IPS = Intra-Parietal Sulcus.

**Table S2.** The locations of the activation peaks (MNI coordinates) from the ICA.

| Functional Network | Cluster size | *T* | x | y | z | Location |
| --- | --- | --- | --- | --- | --- | --- |
| Semantic Language Network (SLN) | 15207 | 33,63144 | -51 | 24 | 12 | L IFG p. Triangularis |
|  |  | 31,55019 | -57 | -15 | -9 | L Middle Temporal Gyrus |
|  |  | 30,99941 | -48 | 27 | -6 | L IFG p. Orbitalis |
|  | 1211 | 18,45376 | -9 | 48 | 36 | L Superior Medial Gyrus |
|  |  | 18,2727 | -6 | 15 | 63 | L Posterior-Medial Frontal |
|  |  | 17,4879 | -3 | 54 | 27 | L Superior Medial Gyrus |
|  | 391 | 18,04999 | 6 | -54 | -42 | R Cerebelum IX |
|  |  | 12,85365 | -9 | -48 | -39 | L Cerebelum IX |
|  |  | 10,96627 | 12 | -39 | -33 | Lobule I IV (Hem) |
|  | 32 | 11,77048 | -9 | -39 | -30 | Lobule I IV (Hem) |
|  | 27 | 9,313284 | 30 | -66 | -60 | R Cerebelum VIII |
|  | 13 | 7,890083 | 18 | -18 | 24 | R Caudate Nucleus |
|  | 10 | 7,162607 | 0 | 24 | 18 | L ACC |
|  | 7 | 6,51627 | 36 | -9 | -12 | R Insula |
|  | 3 | 6,284427 | 66 | 0 | 12 | R Rolandic Operculum |
|  | 4 | 6,134505 | -33 | -69 | -60 | Lobule VIIb (Hem) |
|  | 2 | 5,737201 | -12 | -30 | 15 | Thal: Temporal |
|  | 3 | 5,553001 | 24 | -30 | -21 | R Fusiform Gyrus |
| Anterior Saliience Network (ASN) | 13536 | 40,27691 | 51 | 12 | -3 | R IFG p. Opercularis |
|  |  | 36,2644 | -42 | 6 | 0 | L Insula Lobe |
|  |  | 35,70487 | -39 | 18 | -9 | L Insula Lobe |
|  | 1521 | 22,55642 | -33 | -54 | -33 | L Cerebelum VI |
|  |  | 22,45777 | -12 | -51 | -36 | L Cerebelum IX |
|  |  | 21,92574 | 0 | -51 | -36 | Cerebellar Vermis 9 |
|  | 31 | 10,65318 | 36 | -54 | -33 | R Cerebelum Crus 1 |
|  | 311 | 9,361335 | 51 | -69 | -27 | R Cerebelum Crus 1 |
|  |  | 9,289206 | 36 | -87 | -9 | R Inferior Occipital Gyrus |
|  |  | 8,439513 | 54 | -60 | -30 | R Cerebelum Crus 1 |
|  | 29 | 8,200272 | 51 | -27 | -3 | R Superior Temporal Gyrus |
|  | 22 | 7,268538 | -45 | -6 | -42 | L Inferior Temporal Gyrus |
|  | 13 | 7,225836 | 48 | -6 | -42 | R Inferior Temporal Gyrus |
| Default Mode Network (DMN) | 8142 | 36,92516 | -6 | 45 | 12 | L ACC |
|  |  | 36,17663 | 9 | 57 | 21 | R Superior Medial Gyrus |
|  |  | 33,1425 | 6 | 45 | 6 | R ACC |
|  | 2564 | 20,71007 | 27 | -81 | -33 | R Cerebelum Crus 1 |
|  |  | 15,76761 | 21 | -51 | -15 | R Cerebelum IV-V |
|  |  | 15,37668 | 42 | -63 | -39 | R Cerebelum Crus 1 |
|  | 1017 | 16,73102 | -48 | -63 | 36 | L Angular Gyrus |
|  |  | 13,53661 | -45 | -75 | 3 | L Middle Occipital Gyrus |
|  |  | 12,76854 | -39 | -87 | 6 | L Middle Occipital Gyrus |
|  | 133 | 15,76279 | -30 | -81 | -33 | L Cerebelum Crus 1 |
|  | 1197 | 14,70099 | 54 | 0 | -33 | R Inferior Temporal Gyrus |
|  |  | 14,51339 | 42 | 12 | -39 | R Medial Temporal Pole |
|  |  | 13,4084 | 18 | 21 | 12 | R Caudate Nucleus |
|  | 317 | 13,00971 | -48 | 9 | -39 | L Inferior Temporal Gyrus |
|  |  | 9,865301 | -33 | 15 | -12 | L Insula Lobe |
|  |  | 8,1049 | -33 | 21 | -30 | L Temporal Pole |
|  | 108 | 12,2191 | -66 | -18 | -12 | L Middle Temporal Gyrus |
|  | 46 | 10,20353 | -15 | 18 | 12 | L Caudate Nucleus |
|  | 40 | 9,836002 | -24 | -24 | -15 | Subiculum |
|  |  | 7,072544 | -15 | -30 | -21 | L Cerebelum IV-V |
|  | 82 | 9,320195 | 54 | -57 | 36 | R Angular Gyrus |
|  | 119 | 9,011283 | -54 | -27 | 15 | L Superior Temporal Gyrus |
|  | 33 | 6,978061 | -27 | -60 | -15 | L Fusiform Gyrus |
|  | 7 | 6,922428 | 0 | -72 | 0 | L Lingual Gyrus |
|  | 10 | 6,719633 | -45 | -60 | -42 | L Cerebelum Crus 2 |
|  | 18 | 6,675622 | -36 | 18 | 12 | L IFG p. Triangularis |
|  | 10 | 6,459826 | 21 | -3 | -36 | Entorhinal Cortex |
|  | 3 | 6,312524 | -21 | -60 | -54 | L Cerebelum VIII |
|  | 6 | 5,896214 | -18 | -39 | -3 | L ParaHippocampal Gyrus |
|  | 2 | 5,624026 | -21 | -3 | -36 | L ParaHippocampal Gyrus |
|  | 2 | 5,512121 | -15 | 3 | -12 | L Putamen |
| Left Executive Control Network (LECN) | 7641 | 32,16948 | -42 | 48 | -6 | L Middle Orbital Gyrus |
|  |  | 31,61933 | -33 | -72 | 42 | L Inferior Parietal Lobule |
|  |  | 28,714 | -42 | 30 | 24 | L IFG p. Triangularis |
|  | 2083 | 23,47458 | 12 | -81 | -30 | R Cerebelum Crus 2 |
|  |  | 22,42054 | 39 | -72 | -30 | R Cerebelum Crus 1 |
|  |  | 19,8457 | 36 | -72 | -51 | R Cerebelum VII |
|  | 264 | 16,57392 | -57 | -48 | -9 | L Middle Temporal Gyrus |
|  | 228 | 13,58798 | 39 | -63 | 48 | R Angular Gyrus |
|  | 330 | 12,85706 | -15 | -27 | 63 | L Paracentral Lobule |
|  |  | 8,511838 | 24 | -27 | 63 | R Precentral Gyrus |
|  |  | 8,426131 | 3 | -21 | 60 | R Posterior-Medial Frontal |
|  | 653 | 11,75336 | 30 | 42 | -15 | R Middle Orbital Gyrus |
|  |  | 11,63417 | 48 | 48 | -12 | R IFG p. Orbitalis |
|  |  | 11,59785 | 48 | 36 | 24 | R IFG p. Triangularis |
|  | 141 | 11,02185 | -6 | -99 | 9 | L Superior Occipital Gyrus |
|  |  | 7,91895 | -9 | -93 | 27 | L Cuneus |
|  | 434 | 10,21432 | 48 | -12 | 33 | R Postcentral Gyrus |
|  |  | 10,11133 | 60 | 0 | 12 | R Rolandic Operculum |
|  |  | 10,0633 | 33 | -9 | 3 | R Putamen |
|  | 15 | 9,496819 | 12 | -21 | 0 | R Thalamus |
|  | 13 | 8,315154 | 6 | -60 | -54 | R Cerebellum IX |
|  | 94 | 8,301115 | 15 | -93 | 21 | R Cuneus |
|  |  | 8,156825 | 18 | -99 | 9 | R Cuneus |
|  | 26 | 8,110191 | 33 | 15 | 51 | R Middle Frontal Gyrus |
|  | 5 | 6,135665 | 60 | -45 | -12 | R Inferior Temporal Gyrus |
| Right Executive Control Network (RECN) | 18942 | 34,13781 | 45 | -51 | 48 | R Inferior Parietal Lobule |
|  |  | 29,97357 | 36 | 15 | 54 | R Middle Frontal Gyrus |
|  |  | 27,94745 | 39 | -60 | 45 | R Angular Gyrus |
|  | 481 | 20,97945 | -39 | -60 | 48 | L Angular Gyrus |
|  |  | 5,904872 | -63 | -54 | 30 | L Area PFm (Inferior Parietal Lobule) |
|  | 570 | 18,9823 | -36 | 57 | -6 | L Middle Orbital Gyrus |
|  |  | 11,28266 | -15 | 51 | -15 | L Superior Orbital Gyrus |
|  |  | 5,80178 | -12 | 36 | -9 | L Mid Orbital Gyrus |
|  | 60 | 16,80935 | -9 | 9 | 9 | L Caudate Nucleus |
|  | 156 | 10,84424 | -42 | 18 | 39 | L Middle Frontal Gyrus |
|  |  | 7,010253 | -33 | 15 | 48 | L Middle Frontal Gyrus |
|  | 8 | 7,802741 | -36 | 18 | -3 | L Insula Lobe |
|  | 2 | 5,984267 | -33 | 15 | -21 | L Temporal Pole |
|  | 3 | 5,614158 | -12 | 36 | 15 | L Superior Medial Gyrus |

*Abbreviations:* L = left; R = right; p.= Pars; ACC = Anterior Cingulate Cortex.

